# EnzFinder: a sustainable alternative to chemical synthesis

**DOI:** 10.64898/2026.02.12.705490

**Authors:** Akriti Jain, Nishtha Pandey, Arijit Roy

**Affiliations:** TCS Research (Life Sciences division), Tata Consultancy Services, Hyderabad, 500032, India

**Author notes:** Corresponding author –.

**Keywords:** Biosynthesis, enzyme, green chemistry, biocatalyst, substrate promiscuity

## Abstract

Enzymes have emerged as an important alternative to traditional catalysts in chemical industries over the past few decades owing to their sustainable nature. The application of enzymes in chemical synthesis relies on their ability to catalyse promiscuous reactions. Promiscuous activity of enzymes is an abundant phenomenon in nature; approximately 37% of *Escherichia coli* K12 enzymes show promiscuous activity. This highlights the vast expanse of chemical reactions that can be made biochemically feasible through selection of the correct candidate enzymes. Here, we present EnzFinder, a promiscuous enzyme prediction tool that filters candidate enzymes based on similarity in chemical transformation patterns and subsequently ranks them using substrate–product similarity, enabling enzyme prioritization up to the fourth level of EC classification without requiring sequence information. On a blind benchmarking set of 2,309 biochemical reactions, the method achieves substantially higher prediction accuracy than existing rule-based and deep-learning approaches, with improvements exceeding 20% at the sub-subclass level and significantly higher coverage at the fourth level. Application to industrially relevant reactions demonstrates EnzFinder’s ability to identify alternative enzymes with higher substrate similarity and improved kinetic potential. Furthermore, integration of EnzFinder with in silico retrosynthesis tools enables effective prioritization of enzymatic steps within hybrid chemical–biological pathways. Together, these results establish EnzFinder as a practical and interpretable tool for accelerating enzyme discovery and promoting greener, enzyme-driven synthesis routes.

## Introduction

Enzymes catalyse reactions by accelerating substrate to product conversion. Promiscuous enzymes are the ones that can catalyse multiple reactions. The ability to process multiple substrates, rather than being limited to a specific substrate is known as substrate promiscuity ^1^. It supports the evolution of secondary functions in nature ^2^. Substrate promiscuity can similarly be explored in laboratory to synthesize a wide range of compounds. Enzymes usually act under milder, greener conditions compared to traditional chemical practices. Therefore, they align well with the principles of green chemistry ^3^. The starting point for biosynthesis of a target chemical compound is the identification of suitable enzymes. It involves screening of promiscuous enzymes that can catalyse the desired chemical transformation. Domain experts can draw analogy between chemical and biochemical reactions and suggest promiscuous enzymes for chemical reactions. However, the manual process is time-consuming and may fail to perform an exhaustive search and provide a conclusive solution. Hence, in silico methods that support computer assisted synthesis planning (CASP) is the need of the hour for green chemistry. The success of these methods relies on two important pillars, a comprehensive knowledgebase, and an efficient search approach.

The Enzyme Commission (EC) number is a four-part specification system that provides a standardized, hierarchical classification of enzymes. Enzymes are classified into seven broad classes, viz. oxidoreductase, transferase, hydrolase, lyase, isomerase, ligase, and translocase, based on the type of catalysed reaction. The first digit of EC number specifies one of the seven class. The classes are further divided into subclasses which give more details of the type of reaction and is encoded by the second digit. The third position usually tells about the nature of donors and acceptors while the fourth digit indicates substrate specification ^4^. For instance, the EC number 2.1.1.1 denotes a transferase (class 2) that transfers a one carbon (subclass 2.1) methyl group (2.1.1) to nicotinamide (2.1.1.1). MetaCyc, BRENDA, Rhea, ExplorEnz, KEGG pathway are examples of the knowledgebase that archive details of EC number and their associated reactions ^5–10^. CASP methods process these resources to encode the underlying reaction information in different ways ^11–13^. Rule-based methods summarize the chemical changes that drive reactant to product conversion in the form of transformation patterns, e.g., RDM-patterns and SMARTS strings ^14,15^. AI/ML-based methods, use vector embeddings of the reaction fingerprints as the model feature and treat EC-prediction as a classification problem in general ^16,17^.

One of the early works in this domain used MOLMAP descriptors for encoding the reaction and random forest (RF) model for EC assignment ^18^. ECOH ^19^, another machine learning model, was based on the application of support vector machine (SVM) for enzyme classification. It involved identification of common substructures between substrates and products using maximal common substructure (MCS) search followed by EC assignment using mutual information (MI) scores. Recent advancements in this field include development of transformer architecture based deep learning models. However, data-driven methods have their own limitations, e.g., dataset imbalance significantly limited the prediction accuracy of BEC-Pred method at sub-subclass level for the less represented classes ^20,21^. A multilayer perceptron-based model attempted to address this challenge through explainability framework which involved human-in-the-loop for analysing incorrect classifications and refining model architecture or training data ^22^. Another EC predictor, CLAIRE, approached skewed representation of enzyme class through contrastive learning architecture and data augmentation ^23^.

Contrary to AI/ML-based methods, rule-based methods don’t require large, balanced datasets. These methods use predefined reaction rules to make predictions and offer explainability. E-zyme method uses RDM pattern for characterization of reaction centre and its neighbours ^15^. It was developed more than a decade ago and worked well for reactions with a single reactant-product pair ^24,25^. ECAssigner and EC-BLAST are two reaction similarity search tools that were built around the same time. ECAssigner computed Euclidean distance between reaction diversity fingerprints (RDF) for EC assignment ^26^. EC-BLAST took three metrics, viz. bond-change, reaction centre, and structure similarity into consideration ^27^. Though useful, the methods described above limit the prediction to sub-subclass level specification of enzymes. This leaves the user with a yet to be explored, vast search space.

BridgIT and SelenzymeRF are two state-of-the-art methods that provide details of EC number till the fourth level ^28,29^. BridgIT first recognizes reaction centres through reaction rules screening, followed by reaction fingerprint generation and EC assignment based on a global Tanimoto score. Despite an overall high accuracy, BridgIT has limited prediction capability for unbalanced reactions. In contrast, SelenzymeRF, an updated version of Selenzyme ^30^, performs well with unbalanced reactions. It introduces an enhanced scoring scheme to weight reaction fragments (RF) according to their relevance. However, SelenzymeRF needs host strain details for mapping enzyme sequences to query reactions based on phylogenetic distance. This feature makes SelenzymeRF well suited as a synthetic biology tool but might restrict it from a comprehensive promiscuity search across all enzyme classes.

Here, we present EnzFinder, a rule-based reaction similarity search method that targets enzyme promiscuity rather than canonical EC prediction. It performs preliminary screening of enzyme classes based on similarity in chemical transformation patterns and relevance of match. The top ten scorers at sub-subclass level are then prioritized at the fourth level of enzyme specification. The comparison is expanded beyond reaction centres at this stage to bring in the adjacent context; primary reactant and product molecules of dataset are compared to that of query.

EnzFinder outperformed the accuracy of other contemporary methods by 20% to 30% on a test set of biochemical reactions. It also prioritized promiscuous enzymes for synthesis of industrial chemicals indicating the potential to cut short the design-build-test cycle of enzyme engineering. Application of EnzFinder as a downstream pipeline of retrosynthesis tools ^31,32^ helped propose novel enzyme cascade for arformoterol synthesis. These findings suggest that EnzFinder can serve as a valuable resource to augment CASP methods.

## Experimental

### Data processing

#### Biochemical reaction data curation

MetaCyc, one of the widely used biochemical pathways database ^33,34^, was used as the data resource for this work. MetaCyc version 27(v27) was downloaded from the data downloads link accessed on 14-Aug-2023 ^10^. The csv file had 20511 reactions including orphan reactions that didn’t have any EC number assigned. Orphan reactions and reactions which had EC number mapping only up to subclass level, were excluded. Reactions involving macromolecules as main reactant product pair was also removed before further processing since those were beyond the target application of this work. Phrases like acyl-carrier protein which were not directly involved in reaction were substituted with the character ‘U’ (as suggested by ^35^) in reaction SMILES (Fig. S1). Finally, reaction SMILES were checked for redundancy and only one representative reaction was picked to generate a non-redundant dataset.

#### Assignment of biochemical reaction patterns to the reactions

Multiple biochemical reactions might associate with a common type of chemical transformation e.g., transfer of a methyl group from one molecule to the other. These patterns were characterized and assigned to the reactions of the non-redundant dataset. Firstly, a one-to-one correspondence between the atoms of reactant and product molecules was established by atom-to-atom mapping using GraphormerMapper ^35^. Next, the KCF-convoy package was used to assign KEGG atom types to the constituent atoms of reactant and product molecules ^36,37^. Atom types of the mapped elements between reactant and product molecules were then compared to characterize the nature of biochemical transformation. It was represented using the RDM patterns described earlier (Fig. S2) ^15^. Briefly, R represents the atom types of reaction centre in the reactant and product molecule followed by D representing different and M representing matched neighbour atoms of the reaction centre. The atom types of reactant and product molecules are separated by a hyphen (-); the R, D and M part are delimited by colon (:). Unlike the previous method, both element identity and atom type were compared to determine the D part in this work. In addition to the atom identity, KEGG atom type also captures the surrounding context of the atom based on features like functional group, neighbours, number of bonds etc. The predefined list of 68 atom types was included in this work; the list was further extended to 88 atom types (Supplementary file 1), based on the need to differentiate additional atom types, as experienced during results analysis (refer to results section). These changes made the patterns more informative and specific. The RDM patterns generated were mapped to EC numbers of the parent reactions to create a dataset of unique RDM patterns and their catalysing enzymes.

#### Inclusion of cofactor information with the reaction patterns

Cofactors play a crucial role in driving the thermodynamics of a biochemical reaction, e.g., (S)-lactate to pyruvate conversion can be carried out by both EC-1.1.1.27 (NAD+-lactate dehydrogenase) and EC-1.1.2.3 (L-lactate dehydrogenase (cytochrome)), but the computed Standard Gibbs Free Energy differ significantly (3.76 kcal/mol and -10.86 kcal/mol, respectively) due to difference in the cofactor involved ^10,38,39^. Therefore, cofactor information was retained in the dataset while generating the RDM patterns to accommodate user’s preference for specific cofactor in query reaction. A list of common cofactor pairs with their corresponding EC numbers was prepared (Supplementary file 1).

#### Curation of benchmarking dataset

Since MetaCyc reactions were used to develop the EnzFinder workflow, curated BRENDA reactions archived in the EnzymeMap dataset was used to test the accuracy of EnzFinder ^6,40^. The prediction accuracy was compared to other reaction rules-based methods ^24,28^, which derive rules from KEGG reactions. Consequently, 2309 single step, natural reactions lacking cross-reference to either MetaCyc or KEGG reaction identifiers were used as the blind dataset (supplementary file 1). Two recent deep learning models ^22,23^ were also tested on the same dataset to evaluate the comparative efficiency of the proposed method.

In addition to the 2309 biochemical reactions, 20 industrial reactions (supplementary file 1) were curated from literature for which biocatalysts have been proposed. These novel reactions were used to test the real-world application of EnzFinder.

## Methods

### EnzFinder pipeline development

The patterns dataset described in previous section was used to develop a pipeline for screening promiscuous enzymes for chemical reactions. An outline of the workflow is depicted in fig. 1 and the steps involved are described below.

**Fig. 1:**
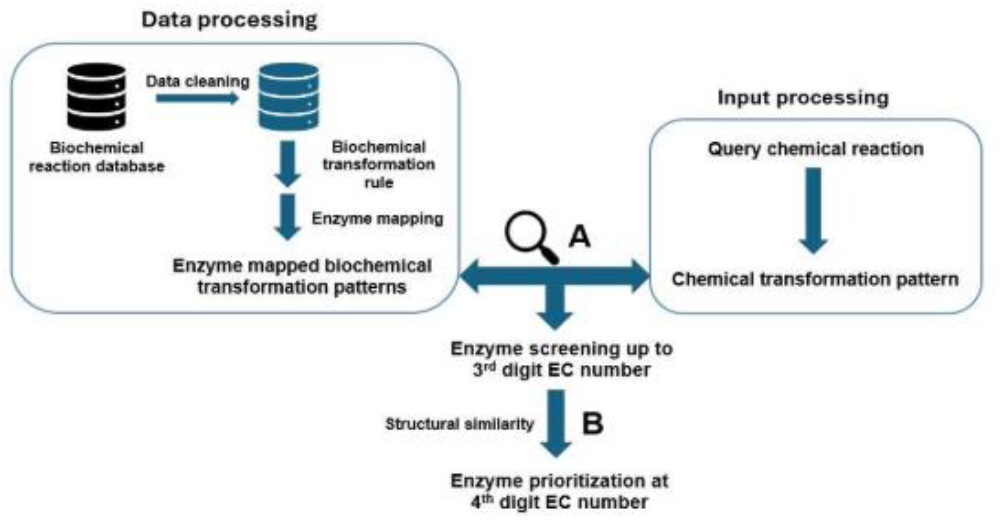
A schematic representation of the EnzFinder workflow

### A) Preliminary enzyme screening to reduce the biochemical search space

Similar to the dataset reactions, atom-to-atom mapping of the query reaction was performed using GraphormerMapper. The RDM pattern for query (RDM_query_) reaction was then generated following the approach described for biochemical reactions. Next, RDM_query_ was scanned against our in-house RDM pattern dataset to explore the biochemical reaction space for similarity. The enzymes corresponding to matched RDM patterns were screened and ranked based on the relative frequency of reported reactions in the nonredundant dataset, using the following expression (equation 1):

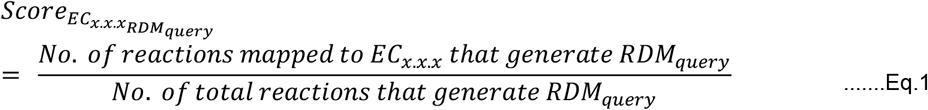

In case of multiple RDM patterns generated from one query reaction, the total score for EC number x.x.x was computed by summing up the individual scores of each matched RDM (equation 2).

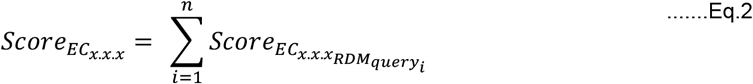

(where n denotes the total number of query RDM patterns that map to EC_x.x.x._)

Considering that an identical transformation might not always be found in the biochemical space, partial pattern matches were also considered to increase sensitivity. The partial patterns considered informative were RD, RpDM, DM, D, RM and R; here RD meant that only the R and D part matched between query and dataset reaction, RpDM meant that in addition to R, M match either the reactant or product D atoms matched with query and so on. This step (step A in Fig. 1) helped in screening EC numbers at sub-subclass level (EC level 3). It reduced the search space of potential enzymes with promiscuous activity. In the next step, (step B, described in the next sub-section) the precise 4-digit EC number was prioritized from the list of enzyme classes filtered in step A. It is important to note that unlike some of the previous methods ^25,41^, this method can process both multi-substrate/product reactions and incomplete reactions without cofactor information as input.

### B) Enzyme prioritization at 4^th^ level of EC number

The steps described above helped in shortlisting a potential set of enzymes based on relevance (Equation 2). However, a higher EC score might not always correspond to a more relevant match due to an uneven distribution of data points across different enzyme classes. This may introduce a bias in the prediction towards the more represented classes. To overcome this bias and to compare the overall structural and topological similarity of query and database reactions, another step was added in the workflow (Fig. 1, step B). The structures of primary reactants and products of top ten EC classes (screened at sub-subclass level described in the previous section A) were compared with the query molecules using the RDKit PatternFingerprint function ^42^. The shortlisted enzymes were re-ranked based on reactant-product pair similarity (Tanimoto scores), neighbouring atoms around reaction centre were considered for sub-subclass level prediction. For prioritisation of the enzyme up to the 4^th^ level details, a dynamic radius quantifying the shortest distance of the farthest atom from the reaction centre was used.

## Results and discussion

### A) Curated dataset of enzyme mapped biochemical reaction transformation pattern

The non-redundant dataset had 13,856 atom-to-atom mapped reaction SMILES associated with 139 unique EC sub-subclasses and 6201 unique EC numbers. As a preliminary quality check, the RDM patterns generated by EnzFinder were manually checked for 250 reaction SMILES. In the process, it was observed that certain atoms could not be classified properly due to lack of a defined atom type e.g., nitrogen atoms in isocyanides (R≡N+-R) or sulfur atoms in sulfoxides (R-S(=O)-R). Hence, the list of KEGG atom types was extended to include wider functional groups (Supplementary file 1); the extended list improved the top-1 accuracy of EnzFinder in preliminary enzyme screening by ∽16%. The subsequent rounds of iterative refinement in the pipeline were thus carried out using the extended list. In total, 88 atom types generated 5932 unique RDM patterns, a summary of the dataset statistics is provided in table 1.

**Table 1.**
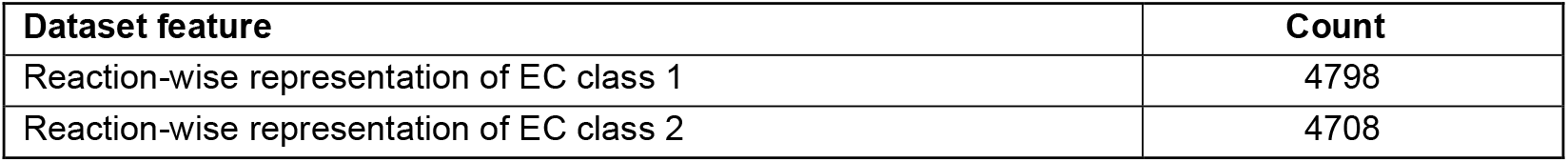
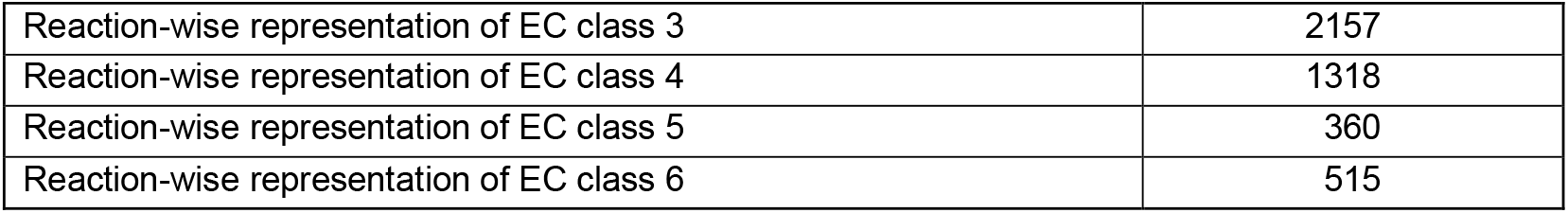
Statistics of the processed dataset (derived from MetaCyc version 27.0)

### B) EnzFinder performance evaluation on biochemical reaction data

The scoring function (equation 2) was designed to pick the most relevant EC numbers from database by assigning weights proportional to the frequency of occurrence. The curated dataset however had a non-uniform representation of different EC class (table 1). This class imbalance was overcome by comparing structures of the substrate-product pairs of the top ten unique EC numbers screened in the first round with that of the query reactions. It captured the degree of similarity between reactions and ensured that the EC number with the most similar substrate-product pair got prioritized. Structure comparison of the dataset and query substrate-product pairs led to reranking of the EC numbers based on Tanimoto score. After reranking, around 20% of the test set reactions got the correct EC at sub-subclass level screened among the highest scorers. The filtering steps reduced the search space for EC prioritization at 4^th^ digit.

The accuracy of EnzFinder was evaluated by checking whether the enzyme mapped to test set reaction was predicted among the top N scorers or not (top-N accuracy). Table 2 summarizes the performance of EnzFinder on the benchmarking dataset (2309 reactions, supplementary file 1) along with other state-of-the-art methods. EnzFinder emerged as the best approach for screening promiscuous enzymes. The top-1 accuracy of EnzFinder on the EnzymeMap test set was significantly better (∼18%-36%) than others.

**Table 2.** Performance evaluation of EnzFinder and other state-of-the art methods on the EnzymeMap benchmarking dataset (2309 reactions) for EC screening up to 3^rd^ digit.

| Method used for EC screening (correct prediction up to 3<sup>rd</sup> digit) | Top 1 prediction matched with test dataset |
| --- | --- |
| E-zyme | 47.81% |
| Theia-RHEA | 65.61% |
| CLAIRE | 57.34% |
| **EnzFinder** | **84.14%** |

The above methods can propose EC number only up to 3^rd^ level (sub-subclass) of specification. Though this helps bring down the search space for testing enzyme activity in an experimental setup, the possibilities to explore remain huge and inconclusive. The statistics summarized in table 1 depicts that 139 EC at 3^rd^ level correspond to over 6000 EC at 4^th^ level, which leaves one with around 45 potential enzymes on an average. Also, it should be noted that multiple enzyme classes can generate the same chemical transformation pattern. Under such conditions, traditionally few enzymes are intuitively picked up from the EC sub-subclass and iterative rounds of activity evaluation is performed through wet-lab experiments ^43^. These efforts can be reduced through methods that predict even the 4^th^ digit of EC classification (table 3).

**Table 3.**
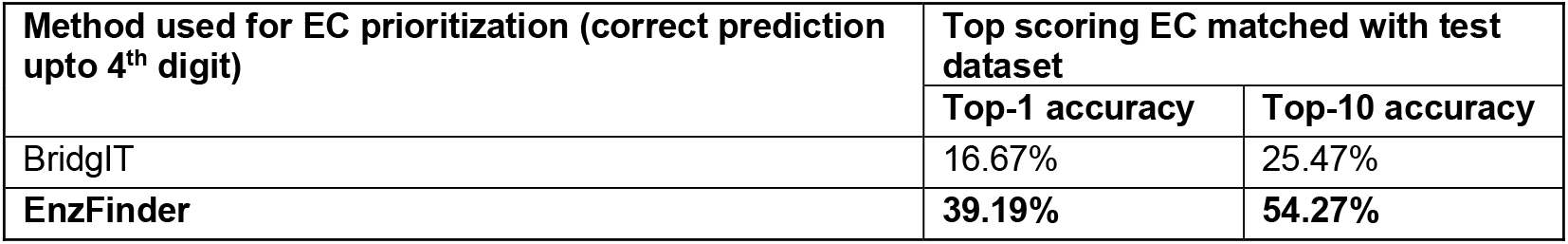
Performance evaluation of EnzFinder and BridgIT on the EnzymeMap test data set for EC prioritization (2309 reactions) up to 4^th^ digit.

| Method used for EC prioritization (correct prediction upto 4<sup>th</sup> digit) | Top scoring EC matched with test dataset |  |
| --- | --- | --- |
|  | Top-1 accuracy | Top-10 accuracy |
| BridgIT | 16.67% | 25.47% |
| EnzFinder | 39.19% | 54.27% |

BridgIT ^28^, another method developed primarily for orphan reaction annotation can predict EC numbers up to 4^th^ digit. Therefore, top-1 and top-10 accuracy of EnzFinder at 4^th^ level of EC number specification were compared with BridgIT (Table 3). As described in the data processing section, 2309 single step, natural reactions lacking cross-reference to either MetaCyc or KEGG reaction identifiers were used as the benchmarking test set. Since BridgIT cannot perform well with incomplete reactions ^29^, only complete biochemical reactions curated in EnzymeMap were used in test set. EnzFinder performed significantly better than BridgIT. BridgIT had significantly lower coverage possibly due to lack of matching rules or incomplete mapping of EC numbers with the rules. It generated output for 58.46% of the reactions from the benchmarking dataset (Table S1 in Supplementary file 1).

The remarkable performance of EnzFinder (table 2-3), was achieved through multiple rounds of refinement. It involved identification of the challenges and their resolution; the following sections describe few relevant examples.

### C) Performance improvement due to expansion of atom types

Atom types define the element identity and surrounding context. A more generic atom type may lead to ambiguity. As mentioned in the data processing section, 21 new atom types were added to the existing 68 atom types described earlier ^15^. Extending the atom list increased the specificity of RDM patterns and led to ∽16% increase in the first round of enzyme screening based on EC score (equation 2). Fig. 2 describes an example where expanding the atom types helped discriminate the reaction centres of enzyme class 1.3.1.x from 1.3.7.x. In the earlier version, both exocyclic (Fig 2a, R03930) and endocyclic double bonds (Fig 2b, R03678) were coded by the atom type C2y (orange). Consequently, KEGG reaction id R03930 (EC-1.3.1.x) and R03678 (EC-1.3.7.x) generated the same combination of R atoms. Adding a new atom type (C2z) for exocyclic double bond differentiated these two reactions. The correct EC (1.3.1.x) was predicted as the top scoring enzyme for EnzymeMap reaction id 74648 with this addition (Fig. 2c).

**Fig. 2:**
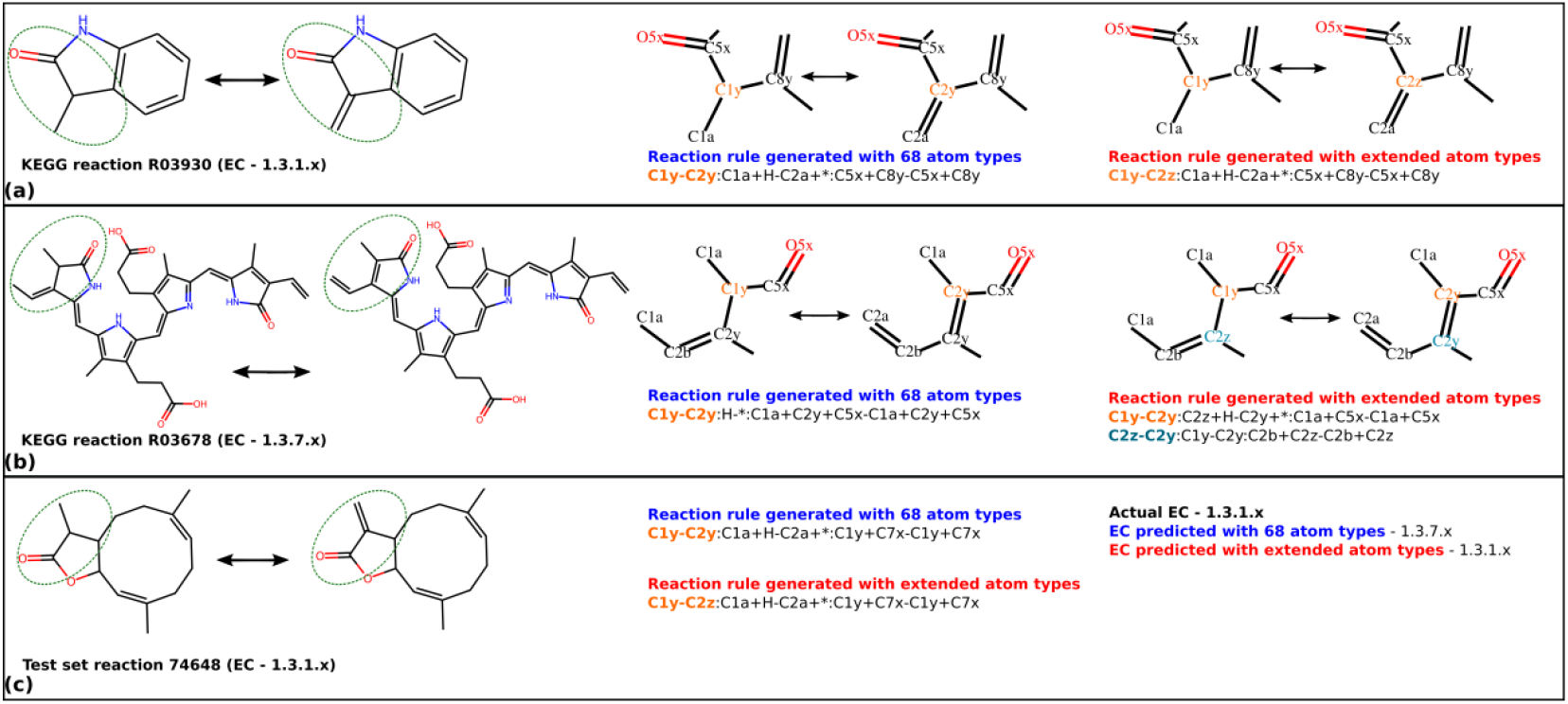
Reaction centres from both (a) EC-1.3.1.x and (b) EC-1.3.7.x generate same pattern with 68 atom types (middle column), but different patterns with additional atom types (right column) introduced in EnzFinder. (c) Query reaction from EC-1.3.1.x was assigned original EC number as the top scorer with the extended atom types.

### D) Inclusion of cofactor to improve specificity

Cofactors regulate the thermodynamics of biochemical reactions, hence the provision to include preferred cofactor information in query reaction was included. Fig. 3a illustrates an example reaction (42601) from the test dataset. The core chemical transformation involved attachment of a hydroxyl group to a carbon atom. A query reaction with only the primary reactant and product picked EC-1.17.4.x as the top scorer, suggesting EC-1.17.4.x possess the ability to catalyse this reaction. Inclusion of the cofactor information prioritized the enzyme class mapped to test set reaction (1.14.11.x) over all others.

**Fig. 3:**
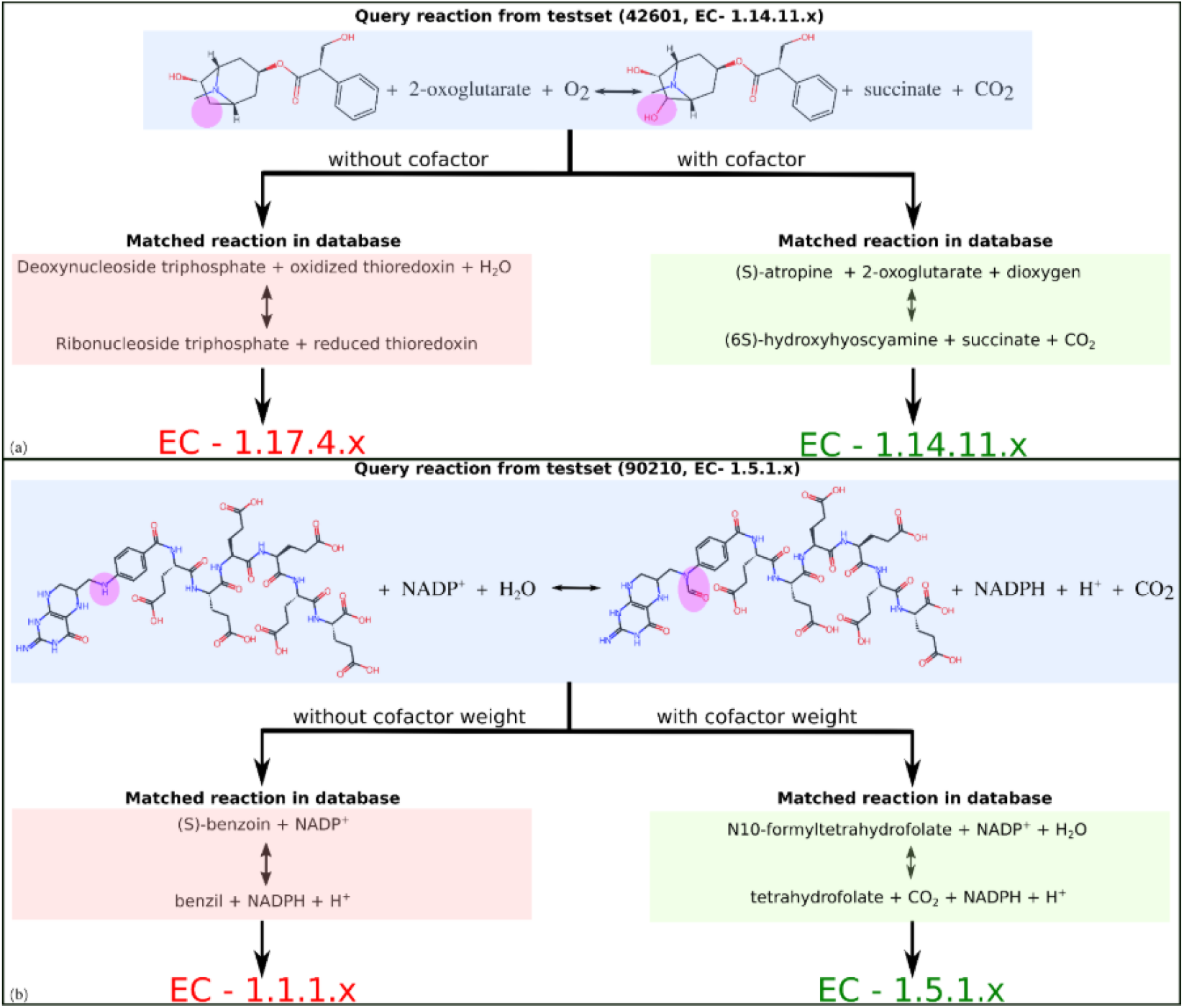
(a) Enzymes from both EC-1.17.4.x and EC-1.14.11.x catalyse attachment of hydroxyl group but differ in the cofactor requirement. Including cofactor information increases specificity of prediction. (b) Different enzyme class share common cofactor, reducing the weight of cofactor match in overall EC score compensated for sparse representation of certain enzymes.

It is important to note that incorporating cofactor information in the query reaction may lead to an adverse outcome due to data imbalance and common cofactor across enzyme classes. Additional care needs to be taken to overcome such scenario. For example, several EC classes are known to share cofactors e.g., most oxidoreductases with EC number 1.x.1.x use NAD+/NADP+ as an acceptor. As highlighted earlier, the data was not uniformly distributed across all EC. In some cases, using cofactor information led to screening of the most abundant EC class (equation 1) with that cofactor, at times superseding the main reactant product similarity. Fig. 3b highlights one such scenario where the test reaction (90210) shared higher similarity with reaction from enzyme class 1.5.1.x (FORMYLTETRAHYDROFOLATE-DEHYDROGENASE-RXN), but EC-1.1.1.x emerged as the top scorer. The cofactor RDM match led to higher reaction count and score of the abundant EC-1.1.1.x (equation 1). This eventually prioritized EC-1.1.1.x over 1.5.1.x, despite dissimilar substrate-product pair. As a corrective measure, the overall contribution of RDM pattern corresponding to a cofactor pair match was reduced in the scoring function. The score in equation 1 was multiplied with a reduced weight (0.5) before adding to the final prediction score. Special attention was given to the consideration that same molecules may serve as substrate for one enzyme and cofactor to other, for instance, acetyl-CoA & CoA pair was considered a cofactor for EC 2.3.1.6 (choline O-acetyltransferase) but not for EC 3.1.2.1 (acetyl-CoA hydrolase). List of the cofactor pairs included in EnzFinder pipeline along with the associated EC number has been provided in Supplementary file 1.

Similarly, it was observed that a less represented EC with full RDM pattern match got a lesser score compared to a more represented EC with partial match (e.g., RM). The levels of matches were thus grouped based on their relevance, e.g., a complete match of R, D and M atom types between query and database reaction pattern was considered more relevant compared to only R or R and M match. Reduced weights were assigned to these groups after optimizing over a range. For example, the highest weight score (1.0) was used for a full RDM match, but lower weights were considered in equation 1 when partial patterns matched (see supplementary file 1, table S2 for details).

### E) Application of EnzFinder as an alternative to chemical synthesis method

EnzFinder was developed with the motivation to propose biochemical alternatives to chemical synthesis methods; 20 reaction SMILES (Supplementary file 1) from the chemical space with known biosynthesis route were tested to evaluate this feature. Table 4 summarizes the top-1 accuracy of EnzFinder and other benchmarking methods up to 3^rd^ level of EC number.

**Table 4.**
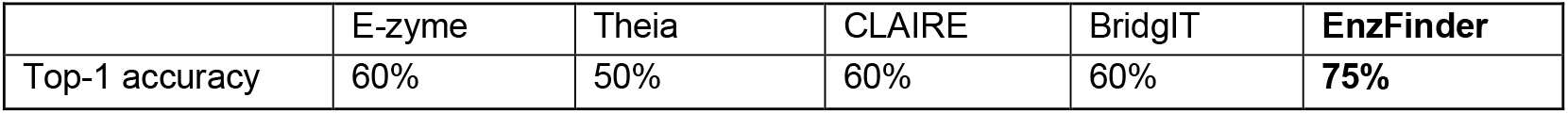
The performance of EnzFinder and other methods on the industrial test cases up to 3^rd^ level of EC number.

In addition to screening the enzymes at sub-subclass level, EnzFinder also proposed potential enzymes with 4^th^ level of specification for 80% of the test cases. Comparison of EnzFinder prioritized enzymatic reaction with the query reaction showed higher similarity in multiple cases; for instance, pantothenate kinase (PanK) is used for substrate phosphorylation in a step of Islatravir synthesis ^43^. EnzFinder scored apulose kinase (2.7.1.233) higher than PanK (2.7.1.33) based on reactant-product similarity. Fig. 4b and 4c compares the natural reactions of PanK and apulose kinase with the industrial reaction (Fig. 4a); apulose kinase reactant-product pair had higher Tanimoto score compared to PanK. Substrate turnover rates of the two enzymes (P0A6I3, Q6D5T8) were also compared using a recent deep learning model CatPred ^44^ to further validate the findings. Apulose kinase showed almost 30-fold higher substrate turnover rate (42.98s^-1^/1.47s^-1^) compared to wild type PanK.

**Fig. 4:**
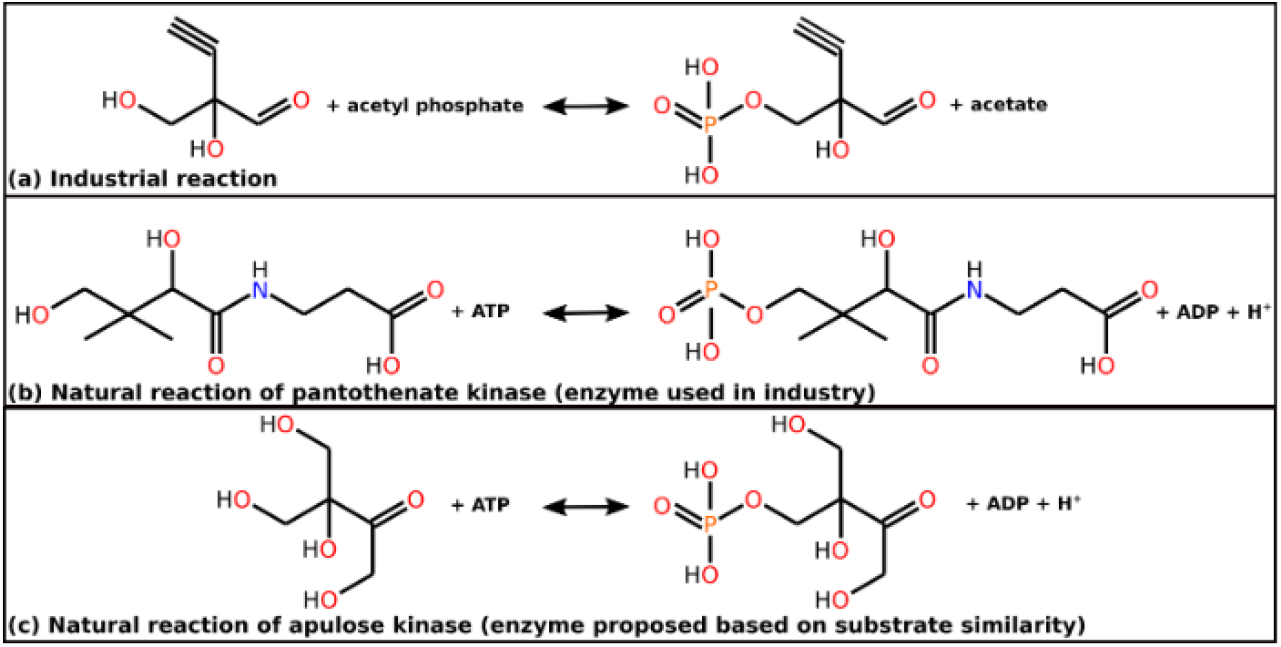
(a) Industrial reaction for Islatravir precursor synthesis compared to natural reactions of (b) PanK (2.7.1.33) and (c) apulose kinase (2.7.1.233). The industrial intermediate shares higher similarity with apulose compared to pantothenate.

Islatravir precursor bound structures of PanK and apulose kinase were further modelled using Boltz2 ^45^ to understand whether it can accommodate the substrate. The predicted structure of apulose kinase and Islatravir precursor complex had > 2-fold higher binding affinity compared to PanK. The substrate binding affinity computed using GNINA ^46^ was -4.43 kcal/mol for wild type apuolse kinase and -2.05 kcal/mol for PanK. Evolution of the enzyme cascade for Islatravir synthesis involved multiple rounds of enzyme engineering and screening ^43^. A pipeline like EnzFinder can significantly reduce the efforts and cost involved in the process.

### F) Augmenting retrosynthesis with EnzFinder

In silico retrosynthesis methods facilitate computer-aided synthesis planning by proposing synthesis routes for a chemical molecule of interest. These methods use reaction templates generated from known chemical or bio-chemical reactions to propose novel synthesis pathways ^11,31,47^. We evaluated the potential of EnzFinder as a tool to augment in silico retrosynthesis. The retrosynthesis route for arformoterol, a bronchodilator prescribed to patients with chronic obstructive pulmonary disease, was built using ASKCOS suite ^48^. It reproduced the hybrid (chemical and enzymatic) route illustrated in one of the recent works by Levin et al. ^31^ (A1-A4 and B1-B3 in Fig. 5). Reaction steps A1, A4, B1, B2 and B3 were proposed as enzymatic reactions. ASKCOS matched the biochemical reaction templates from BKMS-react database ^49^ to the corresponding EC numbers; 2, 6, 2, 31 and 24 unique enzymes were identified as the potential EC numbers for the five enzymatic reactions, respectively. However, lack of prioritization led to multiple equally likely, inconclusive possibilities with difference even at the subclass level. Therefore, there is a need for enzyme prioritization and EnzFinder can play an important role here.

**Fig. 5:**
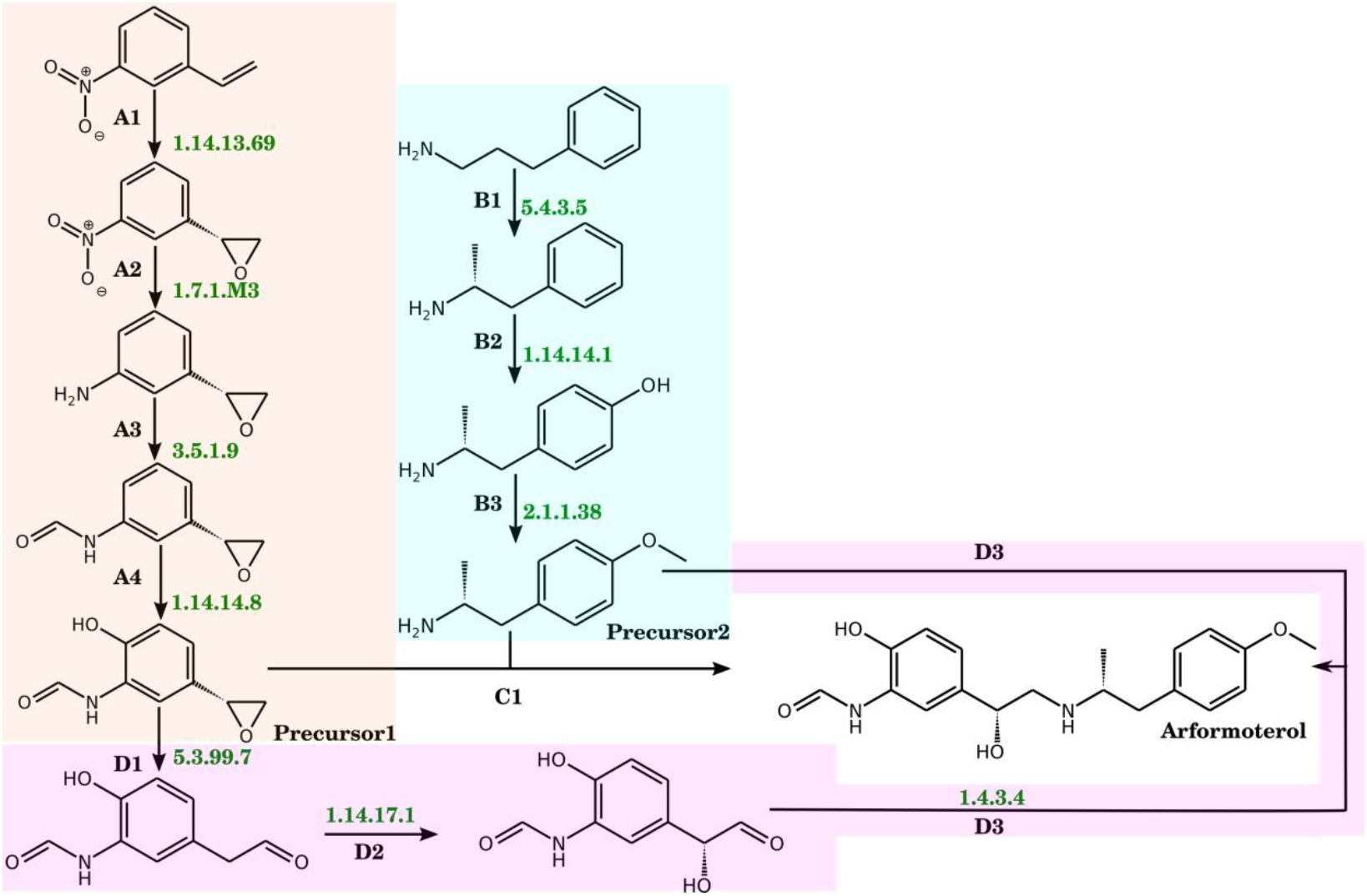
Hybrid retrosynthesis route proposed for arformoterol. Pathways highlighted in peach and cyan suggest set of reactions proposed by ASKCOS ^31,48^ for producing precursor1 and precursor2. Reactions D1-D3 (highlighted in pink) were proposed by DORAnet ^32^ as an enzymatic alternative to C1. EC numbers written against the reactions represent enzymes prioritized by EnzFinder.

The proposed retrosynthesis route of arformoterol involved two parallel pathways for synthesis of precursors (Fig. 5). The two sets of reactions (A1-A4 and B1-B3 in Fig. 5) were queried using EnzFinder. Since EnzFinder checked beyond the reaction centre, it provided a list of EC numbers prioritized according to the degree of substrate similarity. For the proposed enzymatic steps A1, A4, B1, B2 and B3, EnzFinder prioritized EC numbers 1.14.13.69, 1.14.14.8, 5.4.3.5, 1.14.14.1, 2.1.1.38, respectively. Moreover, EnzFinder also predicted EC numbers for chemical reaction steps A2 and A3. Reaction A2 can be catalysed by nitroreductase (1.7.1.M3, RXN-22382) ^10,50^, while A3 can be catalysed by arylformamidase (3.5.1.9, ARYLFORMAMIDASE-RXN).

The final step proposed by ASKCOS was a single step chemical reaction (C1 in fig. 5). To find an alternative biochemical route for this chemical reaction another retrosynthesis tool DORAnet ^32^ was explored. DORAnet proposed a three-step enzymatic reaction (D1-D3 in fig. 5). The reaction rules proposed by DORAnet were next mapped to UniProtKB identifiers ^51^ through which the corresponding EC numbers were identified. In line with the earlier examples, DORAnet also suggested multiple EC numbers that lacked prioritization. EnzFinder provided a prioritized list of EC numbers for steps D1-D3; 5.3.99.7, 1.14.17.1, 1.4.3.4 were proposed for D1, D2 and D3, respectively.

In summary, EnzFinder can complement in silico retrosynthesis/retrobiosynthesis methods by proposing/prioritizing promiscuous enzymes. We hope screening promiscuous enzymes with better substrate resemblance would reduce the rounds of experiments. Additionally, substituting chemical reactions of hybrid route with enzymatic ones would allow development of multi-step enzyme cascades. These cascades promote one-pot synthesis which reduces the operational cost while increasing the overall yield ^52^.

## Conclusion

Enzymes with promiscuous activity can serve as a sustainable alternative to chemical synthesis methods. Usually, promiscuous enzyme screening relies on suggestion from domain experts. In this work, we present EnzFinder, a rule-based, reaction-centric framework that addresses this challenge by enabling automated screening and prioritization of promiscuous enzymes directly from chemical transformation patterns. By combining transformation-level similarity with substrate–product structural comparison, EnzFinder extends enzyme prediction beyond sub-subclass assignment and delivers actionable prioritization at the fourth EC digit, substantially narrowing the experimental search space. Benchmarking against curated biochemical datasets demonstrated that EnzFinder consistently outperforms existing rule-based and machine-learning approaches, particularly for under-represented enzyme classes and incomplete reactions. The incorporation of expanded atom-type definitions and cofactor-aware scoring proved critical for improving specificity while maintaining broad coverage across enzyme classes. Importantly, EnzFinder does not require large balanced training datasets, enzyme sequences, or host information, making it well suited for exploratory searches in enzyme promiscuity and green chemistry applications.

Application to industrial case studies showed that EnzFinder can identify alternative enzymes with higher substrate resemblance and improved kinetic potential compared to enzymes traditionally employed in chemical synthesis. Moreover, coupling EnzFinder with retrosynthesis and retrobiosynthesis tools enabled effective prioritization of enzymatic steps within multi-step hybrid pathways, highlighting its utility as a downstream decision-making layer in computer-aided synthesis planning.

Overall, EnzFinder provides an interpretable and scalable solution for discovering promiscuous biocatalysts and designing enzyme-driven synthesis routes. By reducing reliance on manual expertise and iterative screening, this approach has the potential to shorten design–build–test cycles and facilitate the transition from chemical to enzymatic synthesis. Future integration with enzyme–substrate activity prediction models and thermodynamic feasibility analyses will further strengthen its role in the development of efficient, sustainable biocatalytic pathways.

## Author contributions

Akriti Jain - Data curation, Formal analysis, Investigation, Methodology, Software

Nishtha Pandey - Formal analysis, Investigation, Methodology, Project administration, Validation, Writing – original draft

Arijit Roy – Conceptualization, Methodology, Supervision, Writing – review & editing

## Data availability

The data supporting this article have been included as part of the Supplementary Information.

## Supporting Information

**The Supplementary Information file 1** (MS excel file) contains 5 sheets.

Sheet 1 (S1_CofactorCosubstrateByproduct) - List of cofactor pairs (EC mapped) and common byproducts/cosubstrates/small molecule reactants of biochemical reactions.

Sheet 2 (S2_TestDatasetEnzymeMap) - EnzymeMap reaction identifier, EC number, reaction SMILES of 2309 EnzymeMap reactions used as biochemical reactions test dataset

Sheet 3 (S3_TestDatasetIndustry) - Industrial test cases, Enzyme used in industry, reaction SMILES of 20 industrial reactions used as chemical reactions test dataset

Sheet 4 (S4_atomtype) - List of 88 atom types that were used to characterize the atoms of reactant and product molecules along with their description. New atom types defined in the current work (in addition to the KEGG atom types) are highlighted in bold.

Sheet 5 (S5_Table_S1_S2) - Table S1 summarizes top-1 and top-10 accuracy at 4th digit

**The Supplementary Information file 2** (MS word file) contains 2 figures

Fig. S1: Phrases like acyl-carrier protein are encountered in MetaCyc reaction SMILES, but they are not part of the reaction centre. The phrases were substituted by ‘U’ (MetaCyc reaction RXN-19850). Fig. S2: The atoms of cyclohexanol and cyclohexanone (MetaCyc reaction CYCLOHEXANOL-DEHYDROGENASE-RXN) were assigned their types using the KCF-convoy package. The R, D and M part of RDM patten for the example reaction is highlighted using green, blue and brown colour respectively.

## Acknowledgements

The authors sincerely thank Dr. Rajgopal Srinivasan, Dr. Gopalakrishnan Bulusu, Dr. Navneet Bung, Dr. Broto Chakrabarty, Mrs. Sowmya Krishnan, Mr. Sarveswara Rao Vangala and Ms. Padmasini Raghavachary for their valuable suggestions and comments for the work.

