## Supplementary file 2 for "EnzFinder: a sustainable alternative to chemical synthesis"

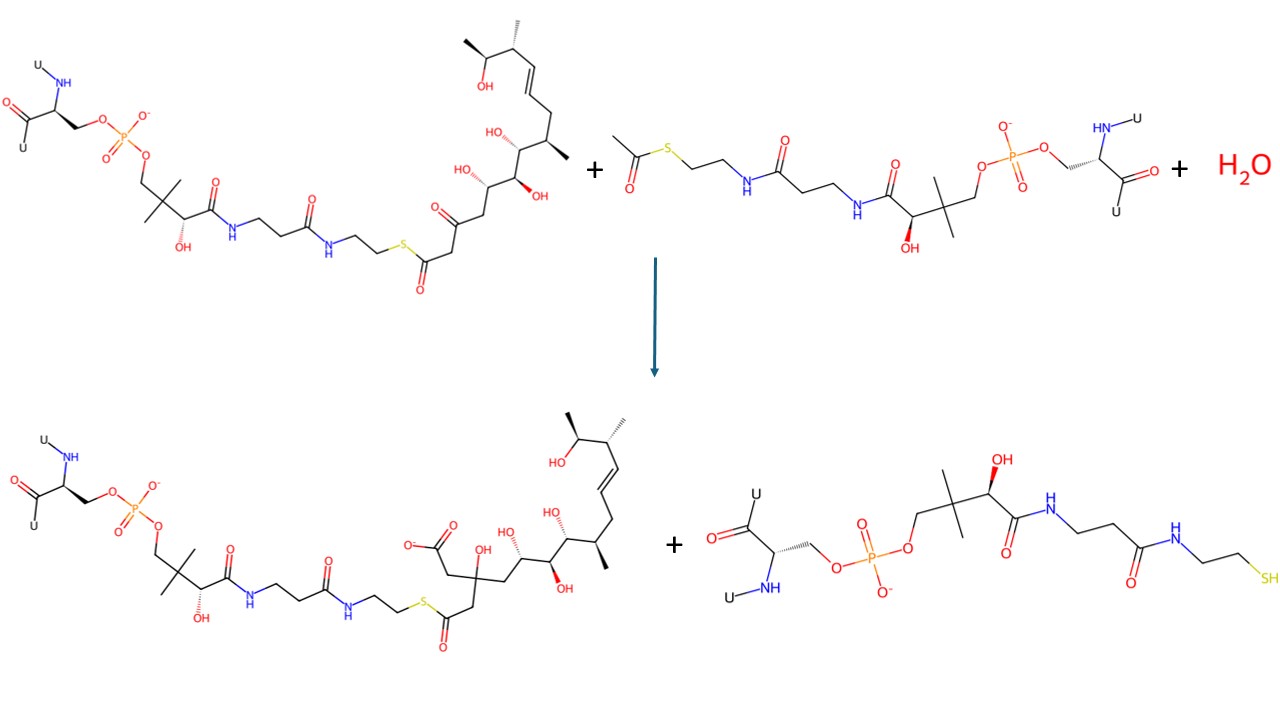

**Fig. S1:** Phrases like acyl-carrier protein are encountered in MetaCyc reaction SMILES, but they are not part of the reaction centre. The phrases were substituted by ‘U’ (MetaCyc reaction RXN-19850).

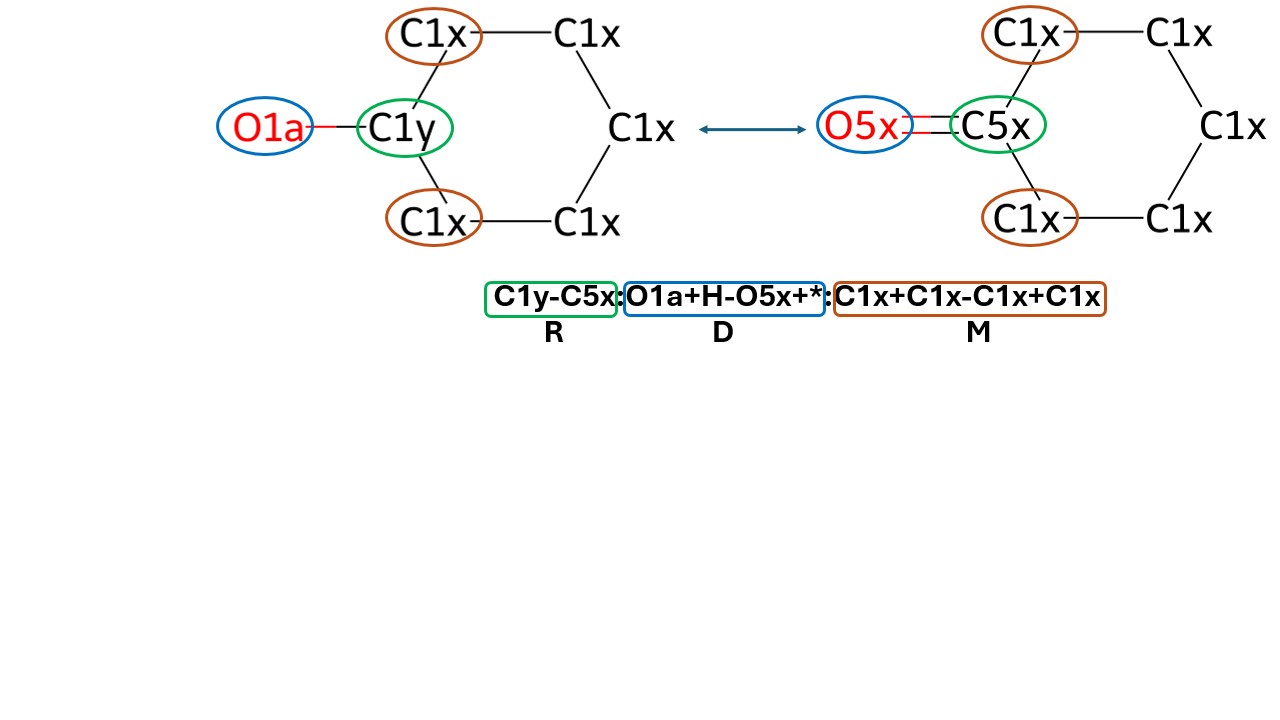

**Fig. S2:** The atoms of cyclohexanol and cyclohexanone (MetaCyc reaction CYCLOHEXANOL-DEHYDROGENASE-RXN) were assigned their types using the KCF-convoy package. The R, D and M part of RDM patten for the example reaction is highlighted using green, blue and brown colour respectively.
